# Cocaine withdrawal enhances subthalamic reward sensitivity

**DOI:** 10.1101/2025.06.10.658581

**Authors:** Maya Williams, Cassandre Vielle, Jeanne-Laure de Peretti, Alix Tiran-Cappello, Christophe Melon, Ali Awada, Yann Pelloux, Nicolas Maurice, Christelle Baunez, Mickaël Degoulet

## Abstract

High-frequency stimulation of the subthalamic nucleus (STN) has shown therapeutic potential in preclinical models of addiction. However, its rewarding properties remain unclear. Here, we show that cocaine withdrawal enhances STN intracranial self-stimulation, revealing a reward-based mechanism that may contribute to the beneficial effects of STN deep brain stimulation in addiction and related neuropsychiatric disorders.

---

Addiction is a chronic psychiatric disorder characterized by a loss of control over dug intake, compulsive seeking and consumption, and repeated cycles of abstinence and relapse ^1^. Preclinical models capturing key diagnostic features, such as dysregulation of intake and persistence of seeking and consummatory behaviors despite negative consequences ^2^, have provided critical insights into the brain mechanisms underlying these behaviors ^3^. Yet, despite extensive research, no pharmacological treatment reliably prevents relapse, particularly in the case of cocaine addiction.

Beyond its well-established role in motor processes, the subthalamic nucleus (STN) has emerged as a key node in cognitive functions, including decision-making or inhibitory control, in both human and rodents ^4–6^. The STN is also the main target for deep brain stimulation (DBS) treatment of both Parkinson’s disease and obsessive-compulsive disorders ^7,8^, and further considered for a wide range of rodent disease models, including addiction ^9^. Manipulations of STN activity, whether via high-frequency (130 Hz) DBS or permanent lesions, consistently reduce cocaine motivation, prevent escalation of intake for multiple drugs, and reduce relapse following protracted abstinence ^10–15^. Moreover, stimulation at lower frequencies modulates compulsive drug seeking in opposite directions: 8 Hz DBS promotes it, whereas 30 Hz reduces it ^16^. Despite these robust behavioral effects, a unifying DBS mechanistic framework is still lacking ^17^. High frequency STN DBS increases dopamine release and metabolism in the striatum of intact and Parkinsonian rats ^18–20^, and further reduces cocaine-induced c-fos expression in striatal regions ^21^, pointing to a potential role for STN in modulating reward-related circuit.

To directly test the rewarding properties of STN activity, we employed an electrical intracranial self-stimulation (eICSS) paradigm in adult male rats. Nose pokes in an active hole, randomly assigned for each individual, triggered was rewarded by a 3-sec electrical stimulation of the STN (50-150µA, 80µs pulse width), while responses in the inactive hole was counted as errors, with no behavioral consequences. Individual stimulation thresholds were determined prior to testing (see methods). Following bilateral stereotaxic implantation of electrodes within the STN, rats (n=11) underwent ten 30-min eICSS sessions at three stimulation (8Hz, 30Hz or 130Hz) ^11,16^, delivered in a pseudo-randomized order (two 5 daily-session blocks mixed between frequencies, Fig. 1A). This design controlled for potential order related to stimulation frequency. Across sessions, naïve rats exhibited a robust frequency-dependent preference for eICSS at 130Hz, with significantly greater responding at 130Hz compared to lower frequencies (Fig. 1B, Repeated One-way ANOVA: *F*_2,18_=29.58, *p*<0.001). eICSS performances remained stable over time, with no significant differences between the first and last five-session blocks for any frequency (Fig. 1C, paired *t*-test: 8Hz: *t*_10_=0.583, *n*.*s*.; 30Hz: *t*_10_=1.69, *n*.*s*.; 130Hz: *t*_10_=1.964, *n*.*s*.). Animals robustly adapted their behavior to frequency-specific stimulation, confirming that 130Hz was the most reinforcing condition (Repeated one-way ANOVA: first block: *F*_2,20_=4.87, *p*=0.019; second block: *F*_2,20_=5.09, *p*=0.016). In support of this, rats performed significantly more perseverative responses (*i*.*e*., unrewarded nose pokes during the post-stimulation 3-sec timeout) at 130Hz than at lower frequencies (Fig. 1D, Repeated One-way ANOVA: *F*_2,18_=9.406, *p*=0.002), while inactive hole activations did not differ across frequencies (Fig. 1E, Repeated one-way ANOVA, *F*_2,18_=2.253, *n*.*s*.). To appreciate frequency-specific reward valuation, we computed a discriminative index (active nose poke / total nose poke), which confirmed strong preference for the active hole at all frequencies (Fig. 1F, paired *t*-test: 8Hz *t*_10_=4.025, *p*<0.001; 30Hz: *t*_10_=4.765, *p*<0.001; 130Hz: *t*_10_=8.316, *p*<0.001), although the index did not differ significantly between frequencies (repeated one-way ANOVA: *F*_2,20_=1.897, *n*.*s*.).

**Fig. 1.**
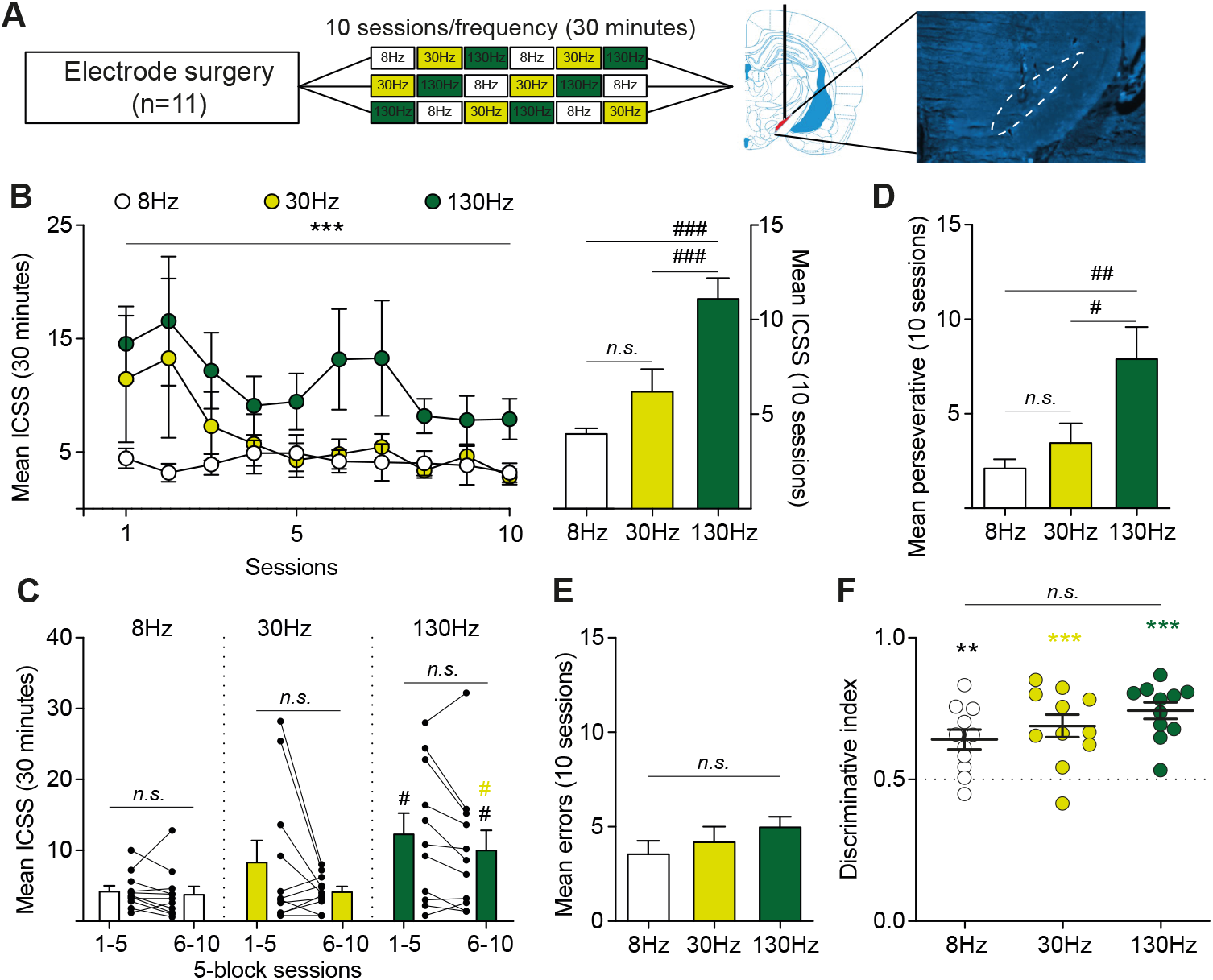
STN neurons support frequency-dependent electrical ICSS. **A**. Experimental design. *Left*: Following bilateral electrode implantation in the STN, naïve rats (n=11) underwent ten 30-min eICSS sessions at 8Hz, 30Hz or 130Hz, pseudo-randomly distributed by blocks of 5-daily sessions. *Right*: Representative histological verification of electrode placement in the STN. **B**. *Left*: Time-course of eICSS responding across the ten sessions. Rats constantly showed a preference for the 130Hz STN stimulation. *Right*: Mean active responding was significantly higher at 130Hz compared to both 8Hz and 30Hz. **C**. eICSS responding remained stable during the initial and last 5 sessions at all frequencies. eICSS at 130Hz was significantly higher than compared to 8Hz (both blocks) and 30Hz (last block). Dots represent individual performance. **D**. Perseverative responses (nose pokes in the active hole during the 3-sec post-reward timeout) were elevated at 130Hz relative to lower frequencies **E**. Error responses (nose pokes in the inactive hole) did not differ across frequency stimulation. **F**. The discriminative index (active pokes/total pokes) was significantly above chance for all frequencies, indicating positive discrimination of STN stimulation. Each dots represents a single subject. Data are presented as mean ± SEM (within-frequency comparisons: ^**^*p*<0.01, ^***^*p*<0.001; between-frequency comparisons: ^#^*p*<0.05, ^##^*p*<0.01, ^###^*p*<0.001).

STN DBS induces both local and widespread effects, distant from the stimulating site ^17^. To determine whether the reinforcing properties of eICSS were locally driven, we used an optogenetic ICSS (oICSS) approach ^22^, enabling selective activation or inhibition of STN glutamatergic neurons, the principal population within the STN ^23^. Animals were bilaterally infected with AAV5 vectors under the CamKIIa promoter to express either the excitatory AAV5-CaMKII-hChR2(E123T/T159C), the inhibitory AAV5-CaMKII-ArchT3.0 or control AAV5-CaMKIIa-EYFP opsins. Optical fibers were implanted above the injection sites, and animals were given the opportunity to self-administer 3-sec light pulses (2ms, 10mW at 130Hz for stimulation; 3-sec pulse, 5mW for inhibition; mix of both patterns for control animals) over ten 30-min oICSS sessions (Fig. 2A). Across sessions, ArchT-expressing rats performed significantly more active nose pokes than either EYFP and hChR2 groups (Fig. 2B, Repeated two-way ANOVA: interaction effect: *F*_18,189_=1.991, p=0.012), suggesting that STN inhibition supports oICSS behaviors. In contrast, oICSS responding in ChR2 rats decreased progressively, indicating a mild aversive response to STN excitation (Fig. 2C, paired *t*-test: EYFP: *t*_9_=0.58, *n*.*s*.; hChR2: *t*_7_=2.889, *p*=0.023; ArchT: *t*_6_=1.93, *n*.*s*.). ArchT animals also showed enhanced responding during the final sessions (two-way ANOVA: *F*_2,13_=11.08, p=0.016), and performed more perseverative responses compared to EYFP control (Fig. 2D, two-way ANOVA: group effect: *F*_2,21_=3.592, *p*=0.046). Inactive responses did not differ across groups (Fig. 2E, two-way ANOVA: group effect: *F*_2,21_=2.788 *n*.*s*.). As expected, EYFP rats displayed a poor discriminative index, indicating their lack of preference between active and inactive holes, ruling out potential light pulse-induced self-administration behavior. ArchT, but not hChR2, animals exhibited a significant elevated discriminative index (Fig. 2F, unpaired *t*-test *vs*. 0.5: EYFP *t*_16_=1.23, *n*.*s*.; hChR2: *t*_14_=1.844, *n*.*s*.; ArchT: *t*_12_=5.558, *p*<0.05), consistent with a rewarding effect of STN neuron inhibition. Finally, in vivo electrophysiological recordings performed in naïve anesthetized ArchT-infected animals confirmed that 3-sec light pulses, delivered every 30-sec, reliably induced rapid, pulse-locked inhibition of STN neurons activity (Fig. 2G), resulting in a transient decrease in STN firing rate (Fig. 2H, Repeated One-way ANOVA: F_2,30_=24.84, p<0.001).

**Figure 2.**
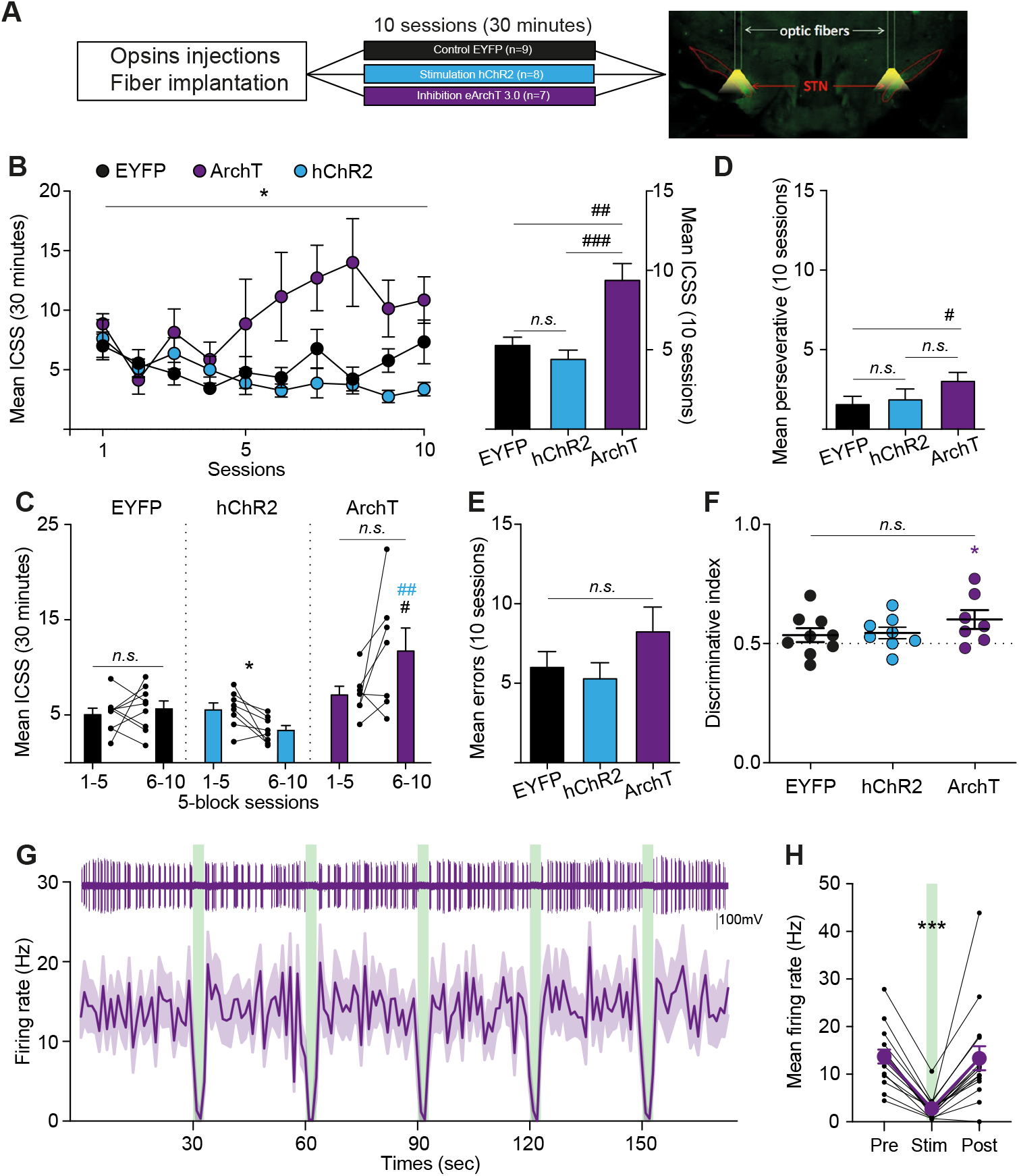
STN optogenetic inhibition supports self-stimulation behaviors. **A**. Experimental design. *Left*: Rats received bilateral STN injection of either AAV5-CaMKIIa-EYFP (control, n=9), AAV5-CaMKII-hChR2-EYFP (activation, n=8) or AAV5-CaMKII-ArchT3.0-EYFP (inhibition, n=7), followed by optic fiber implantation, and underwent ten 30-min oICSS sessions. *Right*: representative histological validation of viral expression and fiber placement within the STN. **B**. *Left*: Time-course of STN oICSS across ten sessions. *Right*: ArchT-expressing rats performed significantly more oICSS reponses than hChR2 and control groups. **C**. oICSS responding decreased over time in hChR2 rats, but remained stable for the control and the ArchT groups. During the final block, ArchT rats showed significantly higher oICSS performance than the other groups. Each dot represents individual animal performance across the first and last blocks. **D**. ArchT rats performed more perseverative responses than control and hChR2 rats. **E**. Error responses did not differ between across groups. **F**. Only the ArchT groups displayed a discriminative index significantly above chance. Dots indicate individual discriminative index. **G**. *Top*: Representative STN unit recording. *Bottom*: In anesthetized ArchT-expressing animals (N=2 rats, n= 16 cells), repeated 3-sec light pulses (green rectangles) induces a rapid and transient inhibition of STN neuronal firing. **H**. Group data showing significant reduction in STN neuron firing rate induced by 3-sec light pulses (n=16 cells, 8-10 light pulses per cell). Data are presented as mean ± SEM (within-frequency comparisons: ^*^*p*<0.05, ^***^*p*<0.001; between-frequency comparisons: ^#^*p*<0.05, ^##^*p*<0.01, ^###^*p*<0.001).

Given that drug withdrawal alters reward sensitivity across multiple brain regions ^24–26^, we next examined whether STN rewarding properties would be affected following cocaine exposure. After fiber implantation and intravenous catheterization, animals (n=12) were trained on a seeking-taking chained schedule of reinforcement to acquire cocaine seeking and taking (250 µg/90 µL per infusion) behavior, and subjected to addiction-like protocols (see methods) to mimic critical features of the human psychiatric diagnosis of addiction ^2,16^. eICSS were performed following a 2-month period of forced withdrawal in their home cage (Fig. 3A). Extended cocaine access (6h per day for 15 days) induced an escalation of cocaine intake over time (Fig. 3B, Repeated two-way ANOVA: session effect *F*_14,140_=5.886, *p*<0.001), independent of future compulsivity status (group effect: *F*_1,12_=0.2168, *n*.*s*.). Compulsive-like seeking behavior was assessed using a punishment contingency paradigm in which animals were required to complete a seeking cycle under probabilistic and unpredictable footshock exposure (0.5mA, 0.5sec, 50% of the completed cycles). Punishment significantly reduced seeking performances across sessions (Fig. 3C, Repeated two-way ANOVA: session effect *F*_7,70_=12.31, *p*<0.001; group effect: *F*_1,10_=10.28, *p*=0.009). Compulsive-like individuals were identified using a composite compulsion score, averaging individual performances during the last four punished sessions ^16^ (Unpaired *t*-test: *t*_10_=4.316, p=0.0015). After a 2-month period of forced abstinence, cocaine-withdrawal rats exhibited a strong frequency-dependent preference for 130Hz stimulation (Fig. 3D, Repeated One-way ANOVA: *F*_2,18_=137.3, *p*<0.001), with a modest increase at 30Hz versus 8Hz. Importantly, eICSS responding did not correlate with compulsivity scores at any frequency (Fig. 3E, linear regression: all *r*^2^<0.25, *n*.*s*.), indicating that eICSS-mediated reward was not modulated by compulsive phenotype. As in naïve animals, eICSS responding remained stable across sessions (Fig. 3F, paired *t*-test: all *t*_11_<1.25, *n*.*s*.) and 130Hz stimulation consistently elicited higher responding across both testing blocks (Repeated One-way ANOVA: first block: *F*_2,22_=14.77, *p*<0.001; second block: *F*_2,22_=11.1, *p*<0.001). Perseverative nose pokes were significantly elevated for both 130Hz and 30Hz stimulation compared to 8Hz (Fig. 3G, Repeated One-way ANOVA: *F*_2,18_=20.42, *p*<0.001), further supporting their rewarding properties. Although error responses increased modestly under 130 Hz (Fig. 3H, Repeated One-way ANOVA: *F*_2,18_=3.604, *p*=0.046), this did not affect the discriminative index, which was significantly above chance for all frequencies and maximized at 130Hz (Fig. 3I, paired *t*-test: all *t*_11_>3.815, *p*<0.01; repeated one-way ANOVA: *F*_2,22_=13.89, *p*<0.001).

**Figure 3.**
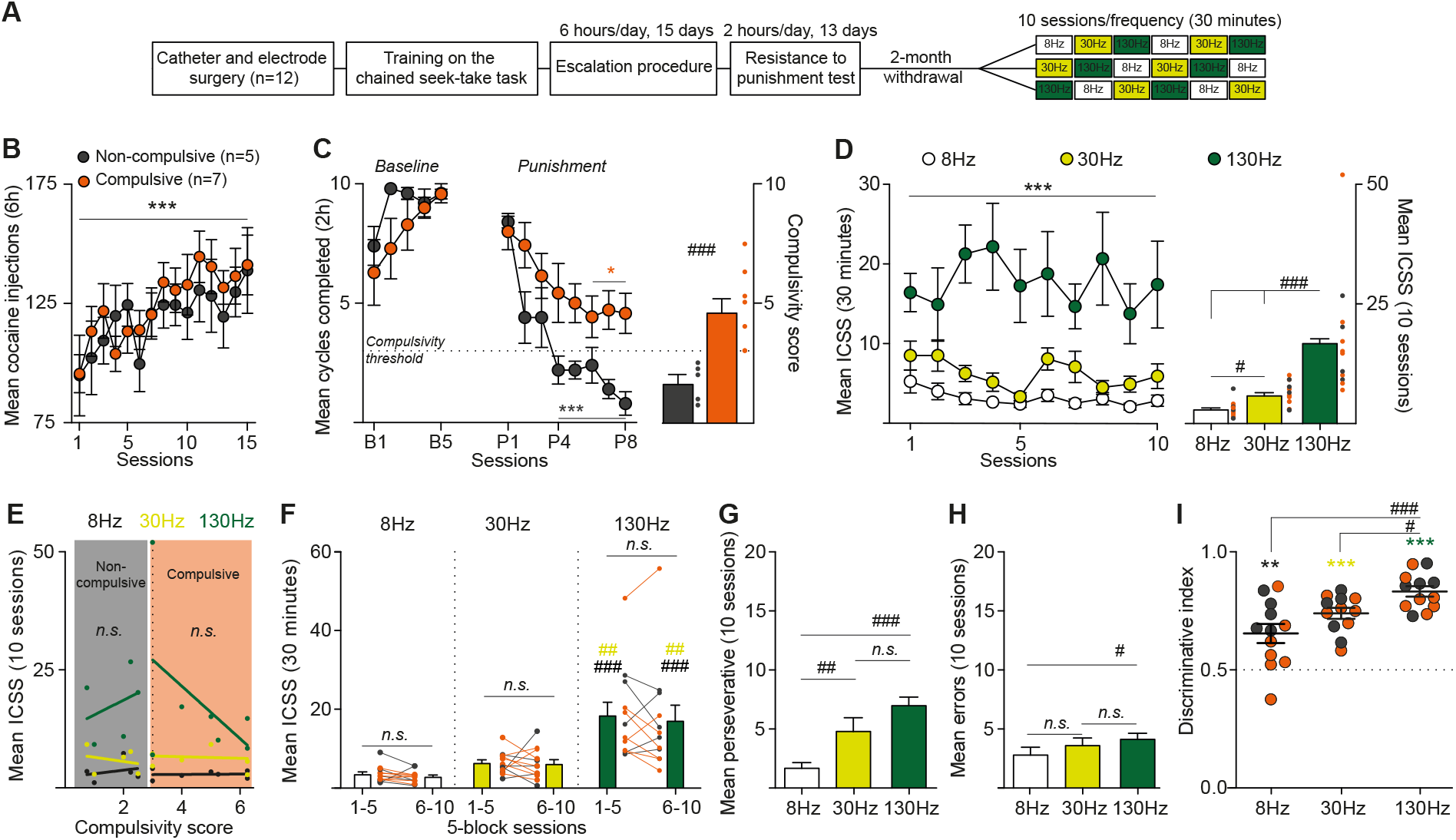
STN electrical self-stimulation is enhanced in cocaine pre-exposed rats after withdrawal. **A**. Experimental design. Rats (n = 12) were implanted with an intravenous catheter and bilateral STN electrodes, trained on a seeking-taking chained cocaine self-administration task, and subjected to an escalation protocol (6h/day for 15 days). Compulsive-like cocaine seeking was assessed under punishment contingency schedule paradigm. After a two-month withdrawal period, animals underwent ten eICSS sessions at 8, 30, and 130 Hz. **B**. Extended access to cocaine led to an escalation of cocaine intake in both *future* Compulsive (n=7) and Non-compulsive (n=5) groups. **C**. *Left*: Punishment contingency revealed compulsive-like individuals, which persisted in seeking cocaine despite foot shocks. *Right*: Compulsivity score, calculated as the mean number of seeking cycle completed over the last four punishment sessions. Dots represent each rats’ score. **D**. *Left*: STN eICSS time-course throughout the ten sessions. *Right*: Rats performed more eICSS at 130Hz compared to both 8Hz and 30Hz. 30Hz responding was also higher than 8Hz. Dots represent each rats’ performance at a given frequency (grey: Non-compulsive, orange: Compulsive). **E**. No correlations were observed between compulsivity score and eICSS performances (Non-compulsive: 8Hz: *r*^2^=0.064, *n*.*s*.; 30Hz: *r*^2^=0.062, *n*.*s*.; 130Hz: *r*^2^=0.107, *n*.*s*.; Compulsive: 8Hz: *r*^2^=0.003, *n*.*s*.; 30Hz: *r*^2^=0.1008, *n*.*s*.; 130Hz: *r*^2^=0.245, *n*.*s*. **F**. eICSS dynamics remained stable across blocks (*t*_11_=1.202, 0.192, and 0.7, for 8Hz, 30Hz and 130Hz, respectively, all *n*.*s*.), with consistently higher responding at 130Hz during both blocks. Dots represents each rats’ performance across blocks. **G**. Animals performed more eICSS Perseverative responses at 130Hz were significantly increased, compared to other frequencies. **H**. Error responses were elevated at 130Hz. **I**. Discriminative index was significantly above chance for all frequencies, with the highest value at 130Hz (8Hz *t*_11_=3.815, *p*=0.003; 30Hz: *t*_11_=10.38, *p*<0.001; 130Hz: *t*_1_=15.17, *p*<0.001). Dots indicate individual discriminative index. Data are presented as mean ± SEM (Within-frequency/phenotype comparisons: ^*^*p*<0.05, ^**^*p*<0.01, ^***^*p*<0.001; Between-frequency/phenotype comparisons: ^#^*p*<0.05, ^##^*p*<0.01, ^###^*p*<0.001).

To assess how cocaine withdrawal alters STN-mediated reward sensitivity, we compared eICSS dynamics in cocaine-withdrawal (n=12) and naïve (n=11) rats across sessions. Nonlinear regression revealed frequency-dependent changes in responding. In naïve rats, eICSS dynamic remained stable at 8Hz (Fig. 4A, *r*^2^= 0.0299), but decreased progressively at 30Hz and 130Hz (*r*^2^=0.802 and 0.5244, respectively; comparison of fits: *F*_4,24_=28.19, *p*<0.001). In contrast, cocaine-withdrawal rats showed diminished responses at 8Hz and 30Hz (*r*^2^= 0.6874 and 0.3211, respectively), while 130Hz dynamic remained constant (*r*^2^=0.00004; comparison of fits: *F*_4,24_=71.98, *p*<0.001), demonstrating a striking divergence from naïve patterns (comparison of fits: 8Hz: *F*_2,16_=8.977, *p*=0.002; 30Hz: *F*_2,16_=3.808, *p*=0.044; 130Hz: *F*_2,16_=15.79, *p*<0.001). Cocaine-withdrawal animals self-administered significantly more 130Hz eICSS than naïve rats (Fig. 4B, Unpaired t-test: *t*_18_=4.809, *p*<0.001), suggesting enhanced reward valuation. Conversely, 8Hz eICSS was reduced in cocaine-withdrawal rats (Unpaired *t*-test: *t*_18_=2.645, *p*<0.05), notably in compulsive-like individuals (One-way ANOVA: *F*_2,27_=4.047, *p*=0.029), with no significant changes at 30Hz (*t*_18_=0.0685, *n*.*s*.). Latency to initiate 8Hz and 30Hz eICSS did not differ between groups and remained unchanged across sessions (Fig. 4C-D, Two-way ANOVA: interaction effect: 1^st^ ICSS: 8Hz: *F*_1,21_=0.59, *n*.*s*.; 30Hz: *F*_1,21_=0.347, *n*.*s*.; 2^nd^ ICSS: 8Hz: *F*_1,10_=3.856, p=0.78; 30Hz:

**Figure 4.**
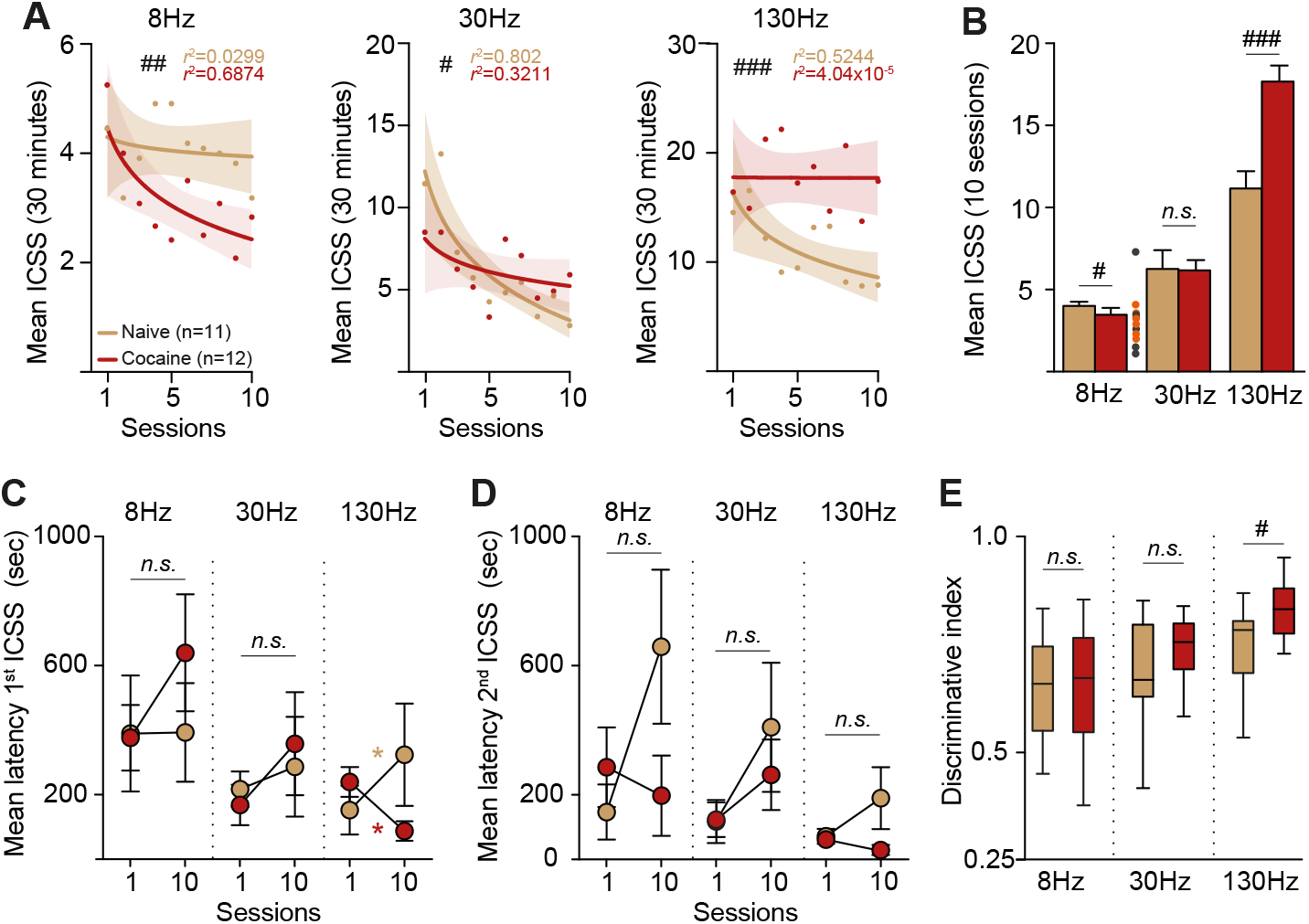
Cocaine withdrawal enhances sensitivity to high-frequency STN self-stimulation. **A**. Nonlinear regression analysis of eICSS behaviors revealed distinct frequency-dependent dynamics between naïve (brown) and cocaine-withdrawal (red) rats. In naïve rats, responding at 8Hz (*left*) remained stable across sessions, but progressively declined at 30Hz (*middle*) and 130Hz (*right*). In contrast, cocaine-withdrawal rats showed reduced responding at 8Hz and 30Hz, while 130Hz performances remained stable over time, indicating a selective enhancement of high-frequency reward sensitivity following withdrawal. **B**. Cocaine pre-exposed animals performed significantly more 130Hz and fewer 8Hz eICSS compared to naïve animals. Dots represents individual performance for compulsive (orange) and non-compulsive (grey) rats. **C**. Latency to initiate the first eICSS remained unchanged at 8Hz and 30Hz in both groups. At 130Hz, latency increased across sessions in naïve rats but decreased in cocaine-withdrawal animals. **D**. Latency to obtain the second eICSS did not differ between groups at any frequency. **E**. Discriminative index was significantly higher for 130Hz stimulation in cocaine-withdrawal rats, with no group differences at 8 Hz or 30 Hz. Each dot represents one subject. Data are presented as mean ± 95% confidence interval (**A**), mean ± SEM (**B, C, D**), and median ± min to max (**E**) (Within-group comparisons: ^*^*p*<0.05; Between-frequency comparisons: ^#^*p*<0.05, ^##^*p*<0.01, ^###^*p*<0.001).

*F*_1,12_=0.5, *n*.*s*.), whereas 130Hz latency diverged: cocaine-withdrawal rats exhibited reduced latency from the first to the last session, while naïve ones showed increased latency (Two-way ANOVA: interaction effect *F*_1,21_=9.839, *p*<0.05). Additionally, cocaine-withdrawal rats tended to press for the second 130Hz eICSS more rapidly than naïve controls in the last session (Two-way ANOVA: interaction effect: *F*_1,19_=3.302, p=0.085). Finally, discriminative index analysis revealed significantly greater preference for the 130Hz stimulation in cocaine-withdrawal rats compared to naïve animals (Unpaired *t*-test: *t*_21_=2.49, *p*=0.021), with no group differences at 8Hz or 30Hz (*t*_21_=0.24 and 1.124, respectively). These results demonstrate that cocaine exposure and withdrawal enhance the sensitivity of STN to high-frequency stimulation, reinforcing its contribution to reward-related behavior.

This study provides direct evidence that STN high frequency stimulation engages reward processes, positioning the STN as a functionally relevant node in the brain reward system. Although the number of self-administered stimulations during STN eICSS sessions was lower than usually observed in well-established reward regions, such as the medial forebrain bundle, ventral tegmental area or prefrontal cortex ^27,28^, both naïve and cocaine-exposed animals reliably discriminated stimulation frequencies. This was reflected in high discriminative index values and a consistent preference for the highest frequency even when tested several days apart. These findings demonstrate that STN stimulation can support reward seeking behaviors. Thus, the level of eICSS responding does not necessarily reflect the reinforcing efficacy of the stimulation site. Mechanistically, our data suggest that 130 Hz STN stimulation drives reward seeking via local neuronal inhibition, as only ArchT-expressing animals exhibited oICSS behavior. Repeated high frequency eICSS likely modulates both afferent and efferent STN pathways ^17^, ultimately leading to dopamine release in reward-related areas ^20^. Cocaine withdrawal is associated with a hypodopaminergic state, which likely contributes to the anhedonic symptoms observed during cocaine withdrawal ^29–31^. In this context, STN DBS may trigger transient phasic dopamine release, compensating for dopamine depletion and sustaining the elevated eICSS behavior seen in cocaine-withdrawal animals. Electrical stimulation-evoked dopamine release in the reward system is further enhanced in the presence of cocaine ^32,33^. A STN DBS-induced dopamine surge could therefore contribute to maintaining dopaminergic tone, sufficient to prevent escalation of cocaine intake and reduce relapse severity for both cocaine and heroin ^11,15^. Together, our results reveal a reward-enhancing dimension of high frequency STN DBS that sustains its beneficial effects on addiction-like behaviors and may further provide a cognitive and reward-related explanation for clinical observations ^34–37^, thereby supporting its use in neurodegenerative and psychiatric disorders.

## Acknowledgments

We thank the National Institute on Drug Abuse Drug Supply Program (NDSP) for generously providing cocaine. We thank the institute’s platforms involved (PAG, NIT, S-prim, UAR 2018). AI-assisted technology was used for spelling and typos correction.

## Funding

Agence National de la Recherche JCJC_ANR-21-CE16-0002 (MD)

NARSAD Young Investigator Grant from the Brain & Behavioral Research Foundation_300015 (MD)

Fondation NRJ_2019 (CB)

Centre National de la Recherche Scientifique and Aix-Marseille Université (CB)

IRESP-19-ADDICTIONS-02 (CB) FRM DPA20140629789 (CB)

Ministry of Higher Education and Research (CV) ENS Lyon (ATC)

Neuroschool of Aix-Marseille Université (MW)

## Author contributions

Conceptualization: YP, CV, CB, MD

Methodology: ATC, YP, NM, CB, MD

Investigation: MW, CV, JLdP, ATC, YP, NM, MD

Visualization: MW, CV, JLdP, NM, MD

Data curation: MW, CV, NM, MD

Funding acquisition: CB, MD

Project administration: CB, MD

Supervision: CB, MD

Writing – original draft: MW, CV, CB, MD

Writing – review & editing: all

## Data and materials availability

All data are available in the main text or the supplementary materials.

## Materials and Methods

### Animals

Sixty four adult Lister Hooded male rats (∼380g, Charles River) used in the experiments were housed in groups of two in Plexiglas cages and maintained on an inverted 12h light/dark cycle (light onset at 7pm) with food and water available ad libitum, in a temperature and humidity-controlled environment. All experiments were conducted during the dark cycle (8am-7pm). All handling and experimental procedures complied with the recommendations for animal experiments issued by the European Commission directives 219/1990, 220/1990, and 2010/63, and approved by Ethic Committee (APAFIS #03129.01, #37070-202204201748941v3, #43676-2023060614092748v4).

### Electrode and Fiber optic implant design

Electrodes were made of Platinum-Iridium wires coated with Teflon (75µm). Coating was removed over 0.5mm at the tips and two wires were inserted into a 16mm long 29G stainless steel tubing to form an electrode. Two of these electrodes, separated by 4.8mm (twice the STN laterally), were soldered to an electric connector, allowing connection with a stimulator. Electrodes were tested with an isolated battery to avoid electrical short circuits. Electrodes (impedance = 20kΩ ± 2.25) and connector were subsequently deeply bound, using a custom mold and dental cement.

Fiber optic implants were built with 230μm optic fibers (NA 0.22, Thorlabs) glued to 2.5mm ceramic ferrules (Thorlabs) using epoxy. Ferrules were slightly grinded with a Dremel to improve the contact with the dental cement.

### Surgery

Animals were anesthetized with either isoflurane (induction 5%/LO_2_ for induction then 2-3%/LO_2_ for maintenance) or a mixture of ketamine (Imalgen, Merial, 100mg/kg, i.p.) and medetominine (Domitor, Janssen, 0.5mg/kg, i.p.), and mounted on a Kopf stereotaxic apparatus (Koft instruments) for bilateral optogenetic viral injections, optic fibers or electrodes implants. Rats received an antibiotic pretreatment by injection of amoxicillin (Citramox, LA, Pfizer, 100mg/kg, s.c.) and an injection of meloxicam (Metacam, Boehringer Ingelheim, 0.5mg/kg, s.c.) for analgesia. After surgery, rats were allowed to recover for at least 7 days with ad libitum access to food and water.

#### Electrical ICSS experiments

Rats (naïve n=22, cocaine-exposed n=12) were implanted with bilateral electrodes within the STN (in mm: -3.7 AP, ±2.4 L from bregma, -8.35 DV from skull surface, with the incisor bar at -3.3 mm ^38^), as previously performed ^11,16^. Four anchoring screws were fixed into the skull. Electrodes, screws, and skull were deeply bounded with dental cement. Cocaine-exposed rats were further a chronically indwelling intravenous catheter. The silicone catheter was inserted and secured into the right jugular vein. The other extremity of the catheter was placed subcutaneously in the mid-scapular region and connected to a guide cannula secured with dental cement. The catheters were daily flushed during the recovery period and just before and after each self-administration session with a saline solution containing heparin (Sanofi, 3 g/l) and gentamicin (Pangram, 4%) to maintain their patency and to prevent infection. Catheters were also regularly tested with propofol (Propovet, Abbott, 10 mg/ml) to confirm their patency. At the end of the recovery period, the stimulation current (50–150µA, 130Hz, unipolar 80µs pulse width) was adjusted for each rats’ STN to the lowest value to observe the hyperkinetic movement in the contralateral paw ^11,13,16^. Stimulation intensity was lowered by 0.1uA and held constant for throughout the experiments.

#### Optical ICSS experiments

Animals (n=30) received a bilateral 0.45μl microinjection of a virus at a rate of 0.16µl/min. To transfect STN neurons, we used AAV5 with the recombinant protein expression under CamKIIa-promoter control (UNC Vector Core, Chapel Hill, USA). Excitatory AAV5-CaMKII-hChR2(E123T/T159C)-p2A-EYFP-WPRE (4×10^12^ pp; n = 11), inhibitory AAV5-CaMKII-ArchT3.0-p2A-EYFP-WPRE (4×10^12^ pp; n=10) or control AAV5-CaMKIIa-EYFP (7.1×10^12^pp; n=9) opsins were injected in the STN, using the same coordinated as above. The injector was left idle at the injection site for 5min to allow efficient diffusion. Optic fibers were implanted 0.4mm above injection site. Electrodes and fiber implants were secured with four anchoring screws and dental cement. The virus was allowed to incubate three weeks before the beginning of optogenetic experiments.

### Behavioral apparatus

All data were acquired on PCs running MED-PC IV or V (MedAssociates).

#### ICSS experiments

Experiments took place rat operant chambers located in sound-attenuating cubicles (MedAssociates). Each chamber was equipped with a house light and two holes with a cue-light. Before each electrical ICSS session, animals were attached to optical fiber connected to a digital stimulation device (DLS8000, WPI) via a stimulus isolator (DLS100, WPI) and a rotating commutator (Doric). For optical ICSS sessions, animals were connected to a 200mW 532nm laser bench (DPSS or Errol laser) controlled by a signal generator (DLS8000_WPI, or Model 3800_A-M System) and connected with an optic coupler (Thorlabs, FCMM625-50A) through a rotary joint (Doric). All ICSS sessions lasted for 30 minutes.

#### Cocaine self-administration experiments

Self-administration apparatus: Behavioral experiments were performed in standard rat operant chambers (MedAssociates), located in sound-attenuating cubicles, equipped with a house light, two retractable levers with a cue light positioned 8 cm above each lever, and a metallic grid floor through which an electric foot shock could be delivered via a generator (MedAssociates). The stainless-steel guide cannula of the catheter was connected through steel-protected Tygon tubing to a swivel (Plastics One), and then an infusion pump delivering cocaine (MedAssociates). Sessions lasted for 2 h or 6 h (see below for detailed procedures).

### Behavioral procedures

#### Electrical ICSS experiments

One week after surgery, or 2 months after the end of the cocaine self-administration protocol, animals were subjected to the eICSS task, consisting in 10 behavioral sessions during which animals could achieve STN self-stimulation at a fixed frequency (8, 30 or 130Hz) for 30 minutes. All animals were subjected to two blocks of 5-daily sessions at a given frequency, several days apart. Order of 5-session blocks were counterbalanced between animals to avoid potential order effect. Stimulation intensity and frequency were fixed for each animal before the ICSS session. The active hole was counterbalanced - right or left - across all animals. Each session started with the illumination of the houselight. Nosepoke in the active hole triggered the delivery a 3-sec electrical stimulation (unipolar pulse, 80µs width) at a given frequency concomitant to a 3-sec illumination of the hole, followed by a 3-sec timeout period, during which nose poke in the active were counted as preservative responses. Nose pokes in the inactive hole had no consequence, and recorded as error responses

#### Optical ICSS experiments

Animals were placed in the same operant conditioning chambers and subjected to the same ICSS paradigm (3-sec light pulse for each active nose poke) than used for eICSS for ten 30-min daily sessions. Stimulation parameters were adapted depending on the group considered (Stimulation: 130Hz, 10mW, 2ms pulse; Inhibition: 5mW, 3s pulse). Before experiments, light power at fiber tip was set, using a power meter (PM20A, Thorlabs), at 10mW for stimulation and 5mW for inhibition. Control EYFP rats were randomly assigned to one light stimulation pattern for the whole experiment.

#### Cocaine addiction-like behaviors

Following catheter implantation and recovery, rats (n=12) were trained (2h per session) to acquire the cocaine taking response, under a fixed ratio 1 (FR1) schedule of reinforcement, for which one press on the active lever (opposite the active one for subsequent ICSS sessions) triggered an i.v. cocaine infusion (250 µg/90 µL over 5 seconds, Drug Supply Program of the National Institute on Drug Abuse). Each cocaine infusion was paired with illumination of the cue light (5 seconds) above the taking lever, retraction of the taking-lever, and extinction of the house light for 5 seconds, followed a 20 second-time out-period (TO).

Animals were then trained on a cocaine seeking/taking-chained procedure ^16^. Each seeking cycle started with the illumination of the house light, insertion of the seeking-lever (2h per session). A single press on the seeking-lever initiated a random interval (RI) schedule of 2 seconds (0.1<RI<4seconds). Seeking-lever presses within the RI (unnecessary seeking lever presses) had no consequences. The first seeking-lever press following the end of the RI completed the seeking cycle, triggering seeking lever retraction, and the insertion of the taking-lever. Press on that lever delivered the cocaine administration, paired with the illumination of the associated cue-light, followed by a 20-second TO, during which both levers were retracted. Following TO, seeking lever was re-inserted to begin the next seek-take cycle. With acquisition of the seeking-taking responses, RI and TO were progressively increase to reach 20, 60, 120 seconds, and 1, 5, 10 minutes, respectively. At the end of training under the seeking-taking schedule, animals were allowed to complete up to 10 seeking cycles during each 2h-session of the RI120-TO10 schedule.

Animals were then subjected to an escalation procedure for 15 consecutive days ^39^, during which they had a daily 6-h period of extended and illimited access to cocaine (FR1). We next tested for their compulsive-like cocaine seeking, using a resistance to punishment test, allowing the identification of *addict* individuals that kept seeking cocaine despite negative consequences ^40^. Following five baseline seeking sessions, as during training (RI120-TO10), rats were subjected to punishment contingency for eight additional 2-h sessions. Here, half of the seeking cycle completed resulted in mild foot shock delivery (0.5 mA, 0.5 s) to the animals’ paws, with no access to the taking lever. Punishment was administered following the first press on the seeking lever after the RI120 schedule has elapsed. The other half of the RI120 completed triggered the insertion of the taking lever, which press initiated cocaine delivery and illumination of the associated cue-light. Seeking cycle’s outcomes (foot-shock or insertion of the cocaine taking lever) were delivered in a pseudorandomized manner, so that difference between both outcomes cannot exceed two. As such, if the two first seeking cycles lead to two consecutive foot shocks, the third cycle would automatically give access to the cocaine-taking lever. Thus, depending on their performances on the punished seeking-taking task, animals could complete up to 10 cycles, leading to a maximum of five foot-shocks and five cocaine infusions. Compulsivity status was extracted from the last four punishment sessions. Animals completing three or more seeking cycles on average were classified as compulsive ^16^. At the end of the punishment test, animals observed a forced-withdrawal period, remaining in their home cage for two months, with no access to cocaine, before being subjected to the ICSS procedure, as described above, for which the rewarded-side has been switched between the two protocols, so that the ICSS nose-poke behavior would not be related to cocaine self-administration.

### in vivo Extracellular inhibition of STN neuronal activity

After at least 3 weeks of recovery following viral injection carrying ARCHT3.0, in vivo electrophysiological recordings were performed in 2 rats) to confirm the ability of STN photoinhibition to suppress neuronal activity. Briefly, rats were anesthetized with a mixture of ketamine/ medetomidine (100/0.5 mg/kg, i.p. supplemented as needed during the recording session) and mounted in a stereotaxic head frame (Horsley-Clarke apparatus; Unimécanique, Epinay-sur-Seine, France). Antalgic treatment comprised metacam (0.5 mg/kg s.c.) and local anesthesia with lidocaine (0.5 ml at 20mg/ml, s.c.). Body temperature was maintained at 36.5°C with a homeothermic blanket controlled by a rectal probe (Harvard Apparatus, Holliston, MA). Single-unit activity of neurons in the STN was recorded extracellularly using glass micropipettes (25-35 MΩ) filled with a 0.5 M sodium chloride solution lowered within the STN. Action potentials were recorded using the active bridge mode of an Axoclamp-2B amplifier (Molecular Devices, San Jose, CA), amplified, and filtered with an AC/DC amplifier (DAM 50; World Precision Instruments). Data were sampled on-line at 10 kHz rate on a computer connected to a CED 1401 interface and off-line analyzed using Spike2 software (Cambridge Electronic Design, Cambridge, UK). An optical fiber (230 μm-Thorlabs) was lowered with a 10° angle just above the STN and connected to a 200mW 532nm DPSS laser. The entry point had the following coordinates: AP: -3.7 mm, ML: +1.0 mm. The fiber tip was at a depth of 7.3 mm from the cortical surface.

The pattern of STN neurons activity was analyzed at least 20s before, 180s during and 20s after photo-modulation. The parameters of light stimulation were the same than those used in behavioral experiments: rats expressing ARCHT3.0 opsin were subjected to ten bins laser stimulation of 3-sec followed by 27-sec OFF (corresponding to 9 consecutive bins for 180 s), with the light power at the tip of optical fiber at 5 mW (±10%).

### Histology and Immunohistochemistry

At the end of the experiments, rats were deeply euthanized with pentobarbital (Dolethal; 200mg/kg i.p.), and perfused with 4% PFA. Brains were removed and flash-frozen in liquid isopentane (−80 °C) to be cut in 40-μm-thick frontal slices with a cryostat in a blinded manner. For electrical ICSS, histological controls for electrodes’ placements were performed after staining with cresyl violet (Fig. 1A). Rat brain slices from optogenetic experiments were processed with immunohistochemistry. Slides were observed under Apotome.2 Microscope (Zeiss) (Figure 1b). Brain slices were washed with PBS 3 times, permeabilized with PBST (0.4% Triton in PBS) for 30min at room temperature, then incubated overnight at 4°C with primary antibody anti-GFP (Life technologies A11120, 1:200). The following day, slices were washed 3 times with PBS and incubated for 2 hours with secondary antibody (Goat anti-mouse IgG, Alexa Flour 488, life technologies A11001, 1:400). Slides were observed under Apotome.2 Microscope (Zeiss, 423667-7144-002). In case of missed electrode, virus, or fiber optic implant placement, the subjects were excluded from final analysis.

A total of 17 animals were discarded from the study after histology. 11 rats were excluded from the analysis for electrode misplacements. A lack of correct expression of GFP was found in 6 rats in the optogenetic groups, which were excluded from the analysis (n=3 in the ArchT3.0 group and n=3 in the hChR2 group

### Statistical Analysis

Data are expressed as mean ± SEM, 95% confidence interval for non-linear regression analyzes (Fig. 4A), or Median and Min-to-Max for whisker analyses (Fig. 4E), with the exact sample size indicated for each group in the figures. Statistics are indicated in figure legends. Data were extracted using a custom-made python pipeline, and analyzed with two-tailed t test, mixed One-or two-ANOVA, mixed-effect models (REML, in case of absence of data, e.g., no 2^nd^ ICSS), and non-linear regression, followed by a Šídák post hoc tests when appropriate, using Prism 10 (GraphPad). Only p-values ≤ 0.05 were considered significant.

